# Adaptation in a fibronectin binding autolysin of *Staphylococcus saprophyticus*

**DOI:** 10.1101/136556

**Authors:** Tatum D. Mortimer, Douglas S. Annis, Mary B. O’Neill, Lindsey L. Bohr, Tracy M. Smith, Hendrik N. Poinar, Deane F. Mosher, Caitlin S. Pepperell

## Abstract

Human-pathogenic bacteria are found in a variety of niches, including free-living, zoonotic, and microbiome environments. Identifying bacterial adaptions that enable invasive disease is an important means of gaining insight into the molecular basis of pathogenesis and understanding pathogen emergence. *Staphylococcus saprophyticus*, a leading cause of urinary tract infections, can be found in the environment, food, animals, and the human microbiome. We identified a selective sweep in the gene encoding the Aas adhesin, a key virulence factor that binds host fibronectin. We hypothesize that the mutation under selection (*aas*_2206A>C) facilitates colonization of the urinary tract, an environment where bacteria are subject to strong shearing forces. The mutation appears to have enabled emergence and expansion of a human pathogenic lineage of *S. saprophyticus*. These results demonstrate the power of evolutionary genomic approaches in discovering the genetic basis of virulence and emphasize the pleiotropy and adaptability of bacteria occupying diverse niches.

**Importance:** *Staphylococcus saprophyticus* is an important cause of urinary tract infections (UTI) in women, which are common, can be severe, and are associated with significant impacts to public health. In addition to being a cause of human UTI, *S. saprophyticus* can be found in the environment, in food, and associated with animals. After discovering that UTI strains of *S. saprophyticus* are for the most part closely related to each other, we sought to determine whether these strains are specially adapted to cause disease in humans. We found evidence suggesting that a mutation in the gene *aas* is advantageous in the context of human infection. We hypothesize that the mutation allows *S. saprophyticus* to survive better in the human urinary tract. These results show how bacteria found in the environment can evolve to cause disease.

## Introduction

Urinary tract infections (UTI) are a global health problem of major significance, with an estimated annual incidence of 150-250 million and a lifetime risk of 50% among women (1–3). The associated costs to individuals and health care systems are substantial, with recent estimates from the United States numbering in the billions per year (4). UTIs are also associated with severe complications such as pyelonephritis, sepsis and premature labor (4). *Staphylococcus saprophyticus* is second only to *Escherischia coli* as a cause of UTI in reproductive aged women (5, 6).

*S. saprophyticus* can be found in diverse niches including the environment, foods, livestock, and as a pathogen and commensal of humans. Several features of the epidemiology of *S. saprophyticus* suggest that infections leading to UTIs are acquired from the environment, rather than as a result of person-to-person transmission (7). This implies that adoption of the pathogenic niche by *S. saprophyticus* has not entailed a trade-off in its ability to live freely in the environment. A recent PCR-based survey of virulence factors in clinical and animal associated isolates showed that *dsdA*, a gene encoding D-serine deaminase that is important for survival in urine (8), and *uafA* and *aas*, adhesins that mediate binding to uroepithelium (9, 10), were present in all isolates surveyed (11), suggesting an underlying pleiotropy, with these virulence factors playing important roles in the diverse environments occupied by *S. saprophyticus*.

The human urinary tract could represent an evolutionary dead end for *S. saprophyticus* with “virulence factors” such as DsdA, UafA and Aas serving an essential function in the primary environmental niche and enabling invasion of the urinary tract as an accidental by-product of this unknown primary function. In this case, we would expect urinary isolates to be interspersed throughout a phylogeny of isolates from the primary niche(s). However, our previous research (7) indicated that human urinary tract infections are associated with a specific lineage of *S. saprophyticus*. Invasion of the human urinary tract enables *S. saprophyticus* to grow to large numbers in urine, isolated from competing bacterial species, before being re-deposited in the environment. This is analogous to *Vibrio cholerae*, which cycles through human and environmental niches and grows to high abundance in the human gut before being deposited in the environment via stool (12, 13). Based on our previous observations and the example provided by other human pathogens that cycle through the environment, we hypothesized that the human urinary tract is an ecologically important niche for *S. saprophyticus* and sought to identify genetic signatures of adaptation to this niche.

The increased availability of sequencing data has enabled comparative genomic approaches that have led to identification of changes in gene content in association with pathogen emergence and shifts in host association. Several notable human pathogens, including *Mycobacterium tuberculosis*, *Yersinia pestis*, and *Francisella tulerensis*, are the product of a single emergence characterized by gene loss and horizontal acquisition of virulence factors (14–16). Similarly, genomic analysis of *Enterococcus faecium* revealed gene gains and losses affecting metabolism and antibiotic resistance in the emergence of a hypermutable hospital-adapted clade that coincided with the profound shift in hospital ecology caused by development of antibiotics (17). Gene gains via recombination have also allowed *Staphylococcus aureus* ST71 to emerge into a bovine-associated niche (18).

Using contemporary and ancient genomic data from strains of *S. saprophyticus*, we found previously that UTI-associated lineages of *S. saprophyticus* were not attendant with specific gene gains or losses; the evolutionary genetic processes underlying *S. saprophyticus’* adoption of the human-pathogenic niche are likely more subtle than what has been described for canonical pathogens (7). Here we have identified one of the mechanisms underlying *S. saprophyticus*’ adaptation to the uropathogenic niche: a selective sweep in the Aas adhesin, which is associated with an apparently large-scale expansion into the human-pathogenic niche. This is, to our knowledge, the first identification of a single nucleotide sweep in a bacterium.

## Results

We reconstructed the phylogeny of *S. saprophyticus* isolates (Table S1) from a whole genome alignment using maximum likelihood inference implemented in RAxML (Figure 1). The bacterial isolates are separated into two clades, which we have previously named Clades P and E (7). In both clades, human associated lineages are nested among isolates from diverse sources, including food (cheese rind, ice cream, meat), indoor and outdoor environments, and animals. Interestingly, cheese rinds harbor diverse strains of *S. saprophyticus*, which cluster with both human- and animal-pathogenic strains.

**Figure 1.**
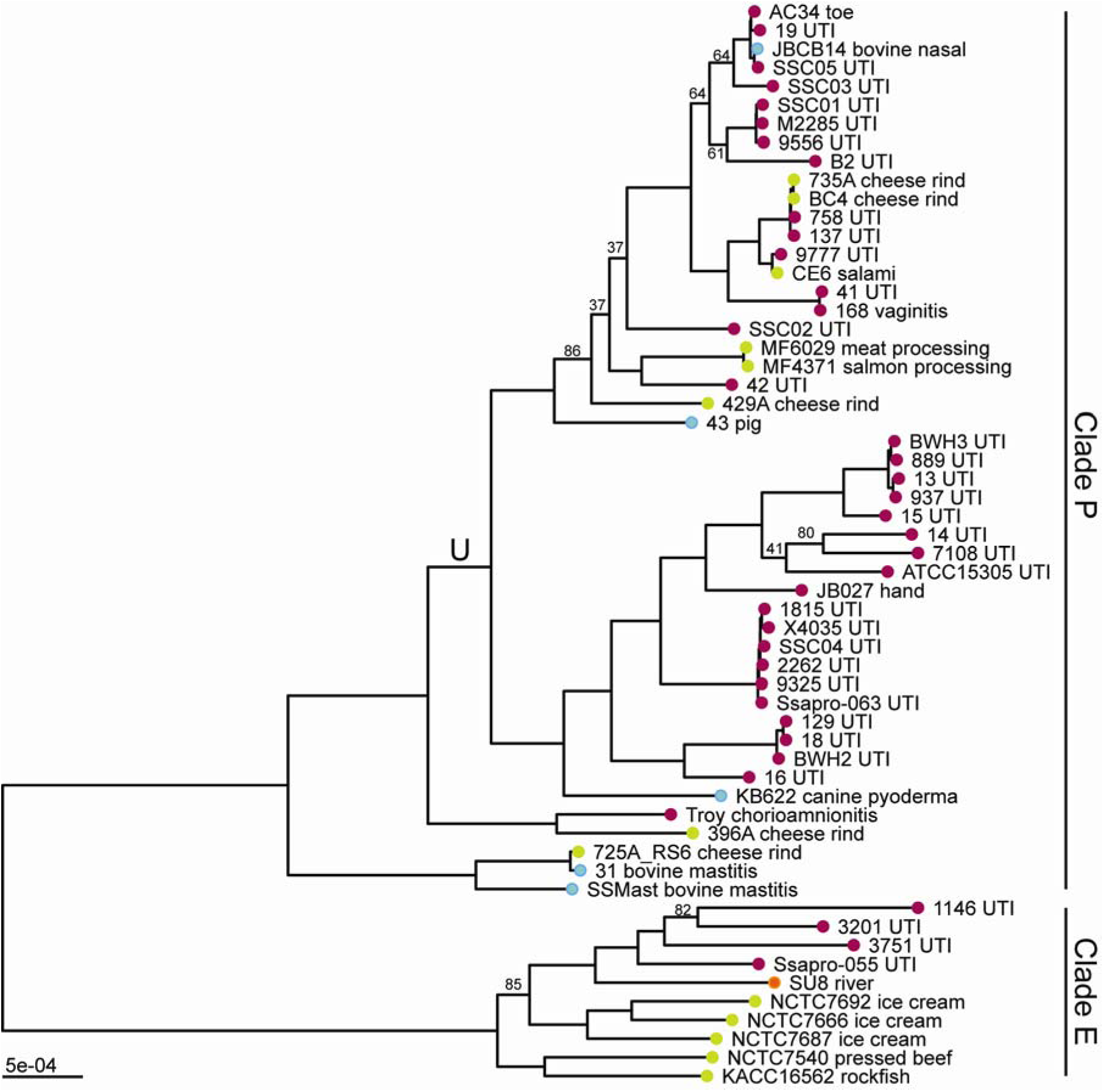
Maximum likelihood phylogeny of *S. saprophyticus*. Maximum likelihood phylogenetic analysis was performed in RAxML (92) using a whole genome alignment with repetitive regions masked. The phylogeny is midpoint rooted, and nodes with bootstrap values less than 90 are labelled. Branch lengths are scaled by substitutions per site. Tips are colored based on the isolation source (pink-human, blue-animal, green-food, orange-environment). Tips are labeled with isolate name and detailed source information. *S. saprophyticus* contains two major clades (Clade P and Clade E). Within Clade P, there is a lineage enriched in human pathogenic isolates (lineage U, branch labeled ‘U’).

Thirty-three of 37 modern, human-pathogenic isolates are found within a single lineage (that we term lineage U, for UTI-associated) to which bovine-pathogenic (mastitis), food-associated isolates, and an ancient genome are basal. Given the association between this lineage and illness in humans, we were curious about its potential adaptation to the human pathogenic niche. The placement of the 800-year old strain between bovine and human-associated lineages suggests it could represent a generalist intermediate between human-adapted and bovid-adapted strains.

Core genome analysis of the 58 isolates of *S. saprophyticus* in our sample showed substantial variability in gene content; the core genome is composed of 1798 genes, and there are an additional 7110 genes in the pan genome. We found previously that uropathogenic isolates of *S. saprophyticus* were not associated with any unique gene content (7). Given the variability in accessory gene content among this larger sample of isolates, we decided to test for relative differences in accessory gene content between human clinical isolates and other isolates using Scoary (19), which performs a genome wide association study (GWAS) using gene presence and absence. We did not identify any genes that were significantly associated with the human pathogenic niche after correction for multiple hypothesis testing using the Bonferroni method.

In addition to the variability observed in gene content, analyses of the core genome also indicated relatively frequent recombination among *S. saprophyticus* (Figure 3). We identified recombinant regions with Gubbins (20), which identifies regions with high densities of substitutions. These results indicated that 70% of sites in the *S. saprophyticus* alignment have been affected by recombination. Recombination can affect bacterial evolution both by introducing novel polymorphisms from outside the population and by reshuffling alleles without increasing overall diversity. Considering sites that are reshuffled within the *S. saprophyticus* sample as recombinant, we estimate a ratio of recombinant to non-recombinant SNPs of 3.4. When considering only the SNPs that introduce novel diversity as recombinant, our estimate of the ratio of recombinant SNPs to non-recombinant SNPs is 0.51. The mean r/m of branches in the phylogeny is 0.82 as estimated by Gubbins (range: 0-7.6). Removal of recombinant SNPs did not affect the topology of the maximum likelihood phylogeny. We observed regional patterns in the amount of recombination inferred, and, as expected, recombination appears frequent at mobile elements such as the staphylococcal cassette chromosomes (SCC_15305RM_ and SCC_15305cap_) and *v*Ss15305 (10).

Adaptation to a new environment may be facilitated by advantageous mutations that quickly rise in frequency, leaving a characteristic genomic imprint: reduced diversity at the target locus and nearby linked loci (i.e. selective sweep, (21, 22)). In order for positive selection to be evident as a local reduction in diversity, there must be sufficient recombination that the target locus is unlinked from the rest of the genome; for this reason, scans for sweeps have been used primarily for sexually reproducing organisms (23–26). As described above and in prior work (7), we found evidence of frequent recombination among *S. saprophyticus*. We hypothesized that *S. saprophyticus*’ transition to the uropathogenic niche may have been driven by selection for one or more mutations that were advantageous in the new environment, and that levels of recombination have been sufficient to preserve the signature of a selective sweep at loci under positive selection. We therefore used a sliding window analysis of diversity along the *S. saprophyticus* alignment as an initial screen for positive selection. We identified a marked regional decrease in nucleotide diversity (π) and Tajima’s D (TD) that is specific to lineage U (Figure 2); TD for this window was -0.38 and 0.94 for non-lineage U Clade P isolates and Clade E isolates, respectively. The region with decreased π/TD corresponds to 1,760,000-1,820,000 bp in *S. saprophyticus* ATCC 15305 and has the lowest values of π and TD in the entire alignment. We investigated the sensitivity of our sliding window analyses to sampling by randomly subsampling lineage U isolates to the same size as Clade E (n = 10); we found the results to be robust to changes in sampling scheme and size.

**Figure 2.**
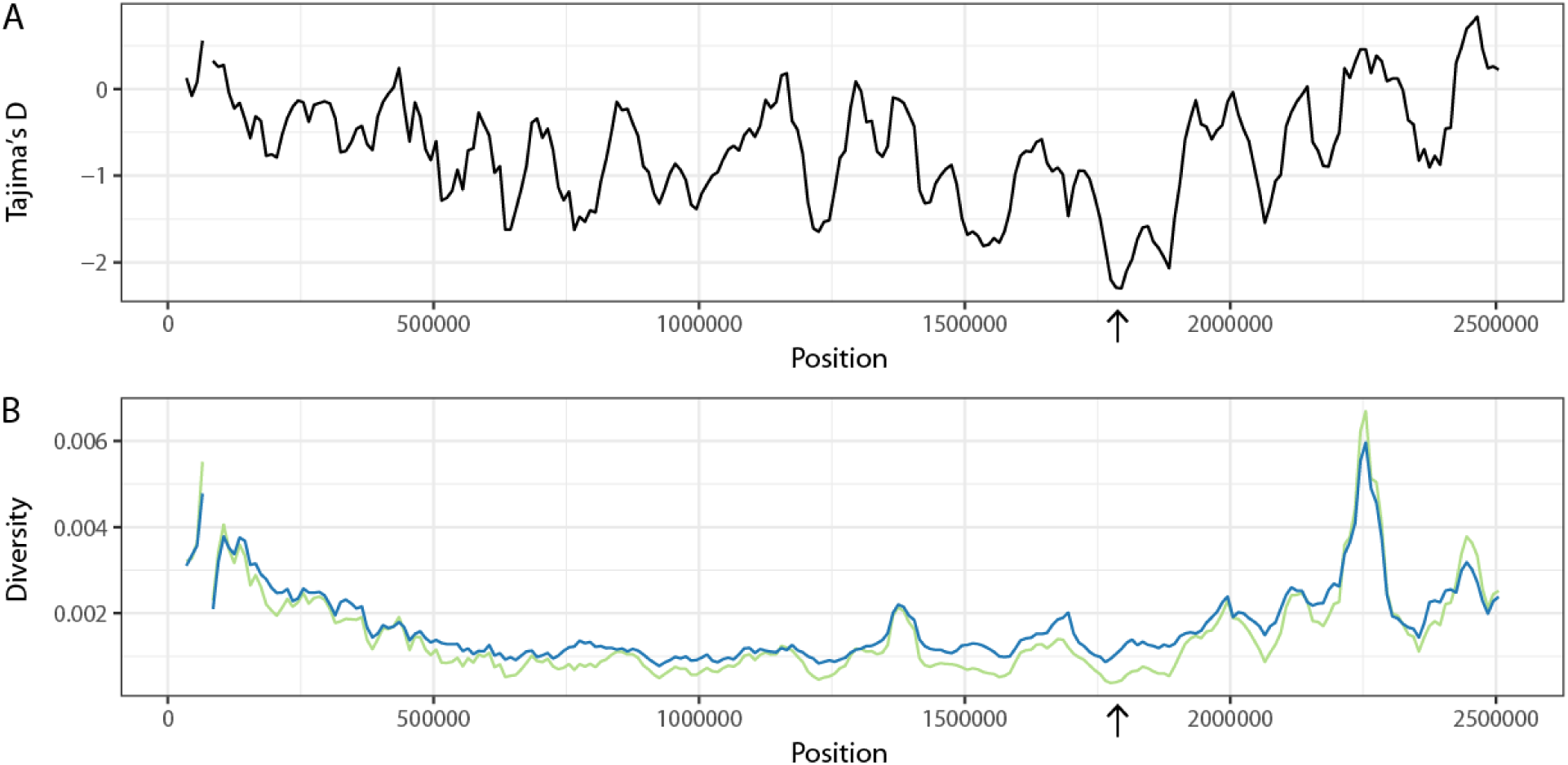
Sliding window analysis of diversity and neutrality statistics. Population genetic statistics were calculated for lineage U using EggLib (94). Windows were 50 kb in width with a step size of 10 kb. A) Tajima’s D. B) π (green) and θ (blue). The lowest values for Tajima’s D and π are found in the same window (1,760,000-1,820,000 bp, arrow).

**Figure 3.**
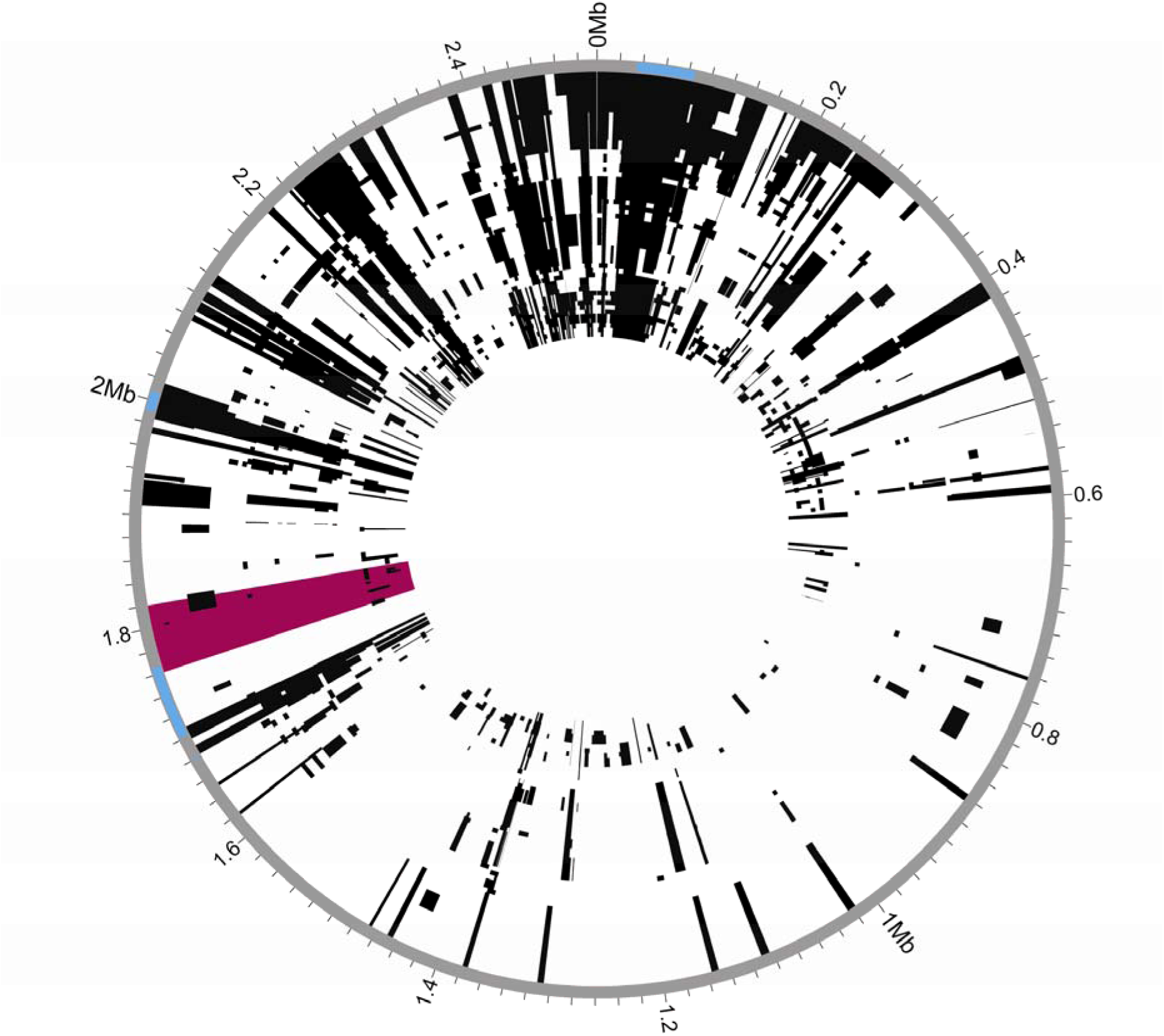
Recombination in *S. saprophyticus*. Recombinant regions in the whole genome alignment of *S. saprophyticus* were identified using Gubbins (20). Mobile genetic elements are highlighted in blue on the outer rim. The window with low Tajima’s D and π is highlighted in pink. Few recombination events are inferred within this region.

To complement the sliding window analysis and pinpoint candidate variants under positive selection, we used an approach based on allele frequency differences between bacterial isolates from different niches. We calculated Weir and Cockerham’s F_ST_ (27) for single nucleotide polymorphisms (SNPs) in the *S. saprophyticus* genome using human association and non-human association to define populations. The region of low π/TD included three non-synonymous variants in the top 0.05% of F_ST_ values (Table 1).

**Table 1.** Single nucleotide polymorphisms with *F*_*ST*_ values in the top 0.05% between 1,760,000 and 1,820,000 bp in ATCC 15305.

| Position | Frequency in human associated isolates | Frequency in non-human associated isolates | $F_{ST}$ Value | Type |
| --- | --- | --- | --- | --- |
| 1772616 | 0.72 | 0.16 | 0.5 | Non-synonymous |
| 1797190 | 0.9 | 0.37 | 0.48 | Synonymous |
| 1808274 | 0.8 | 0.21 | 0.52 | Synonymous |
| 1811585 | 0.72 | 0.16 | 0.5 | Synonymous |
| 1811777 | 0.9 | 0.37 | 0.48 | Non-synonymous |
| 1813204 | 0.74 | 0.16 | 0.53 | Synonymous |
| 1816895 | 0.77 | 0.05 | 0.71 | Non-synonymous |
| 1818150 | 0.77 | 0.16 | 0.56 | Intergenic |
| 1818151 | 0.77 | 0.16 | 0.56 | Intergenic |
| 1818156 | 0.77 | 0.16 | 0.56 | Intergenic |

One of these variants was fixed among human-associated isolates in lineage U (position 1811777 in ATCC 15305, F_ST_ = 0.48) and distinct from the ancestral allele found in basal lineages of Clade P, including the ancient strain of *S. saprophyticus* Troy. This suggests that the variant may have been important in adaptation to the human urinary tract. To assess the significance of the F_ST_ value for this variant, we performed permutations by randomly assigning isolates as human associated, and we did not achieve F_ST_ values higher than 0.28 in 100 permutations.

Selective sweeps may be evident as a longer than expected haplotype block, since neutral variants linked to the adaptive mutation will also sweep to high frequencies (28). Given the evidence suggesting there was a selective sweep at this locus, we used haplotype based statistics to test for such a signature in the *S. saprophyticus* alignment. Haplotype-based methods are hypothesized to not be applicable to bacteria due to differences between crossing over and bacterial patterns of recombination (29), but the methods had not been tested in a scenario akin to a classical sweep, in which local changes in diversity and the SFS have been observed. We found that the variant at position 181177 did show a signature of a sweep using the extended haplotype homozygosity (EHH) statistic (28) (Figure 4). However, the variant did not have an extreme value of *nS*_L_, which compares haplotype homozygosity for ancestral and derived alleles (30), after normalization by the allele frequency.

**Figure 4.**
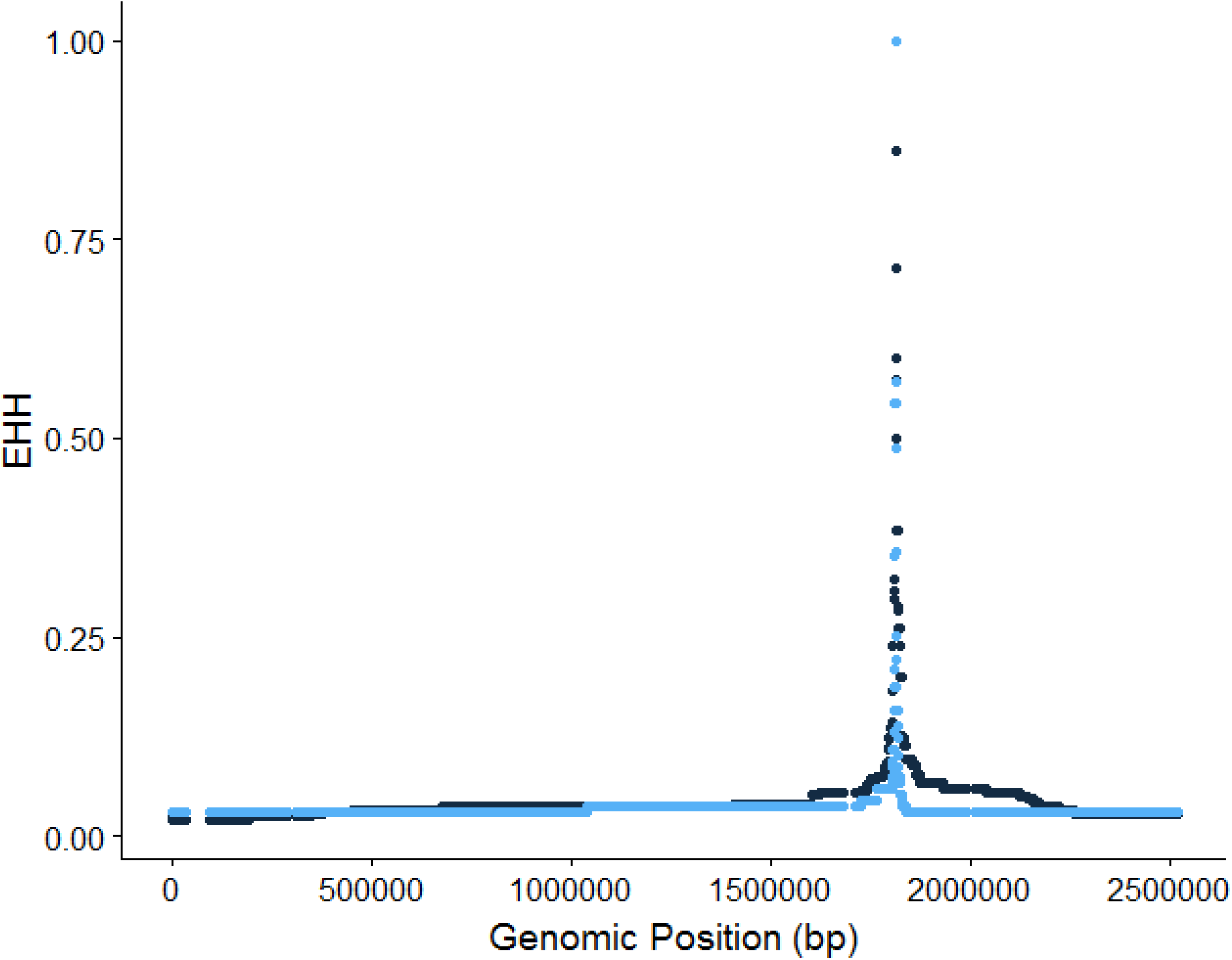
Extended Haplotype Homozygosity (EHH) of single nucleotide polymorphism at position 1811777. EHH values for the ancestral allele are in light blue. EHH values for the derived allele are in dark blue.

The variant of interest (*aas*_2206A>C) causes a threonine to proline change in the amino acid sequence of Aas, a bifunctional autolysin with a fibronectin binding domain (Figure 5, (31)). There are 8 additional nonsynonymous polymorphisms in the fibronectin binding domain; however, none are as highly associated with human pathogenic isolates (Table 1). Adhesins such as Aas are important in the pathogenesis of *S. saprophyticus* urinary tract infections, and this gene has been previously implicated as a virulence factor (9, 31, 32).

**Figure 5.**
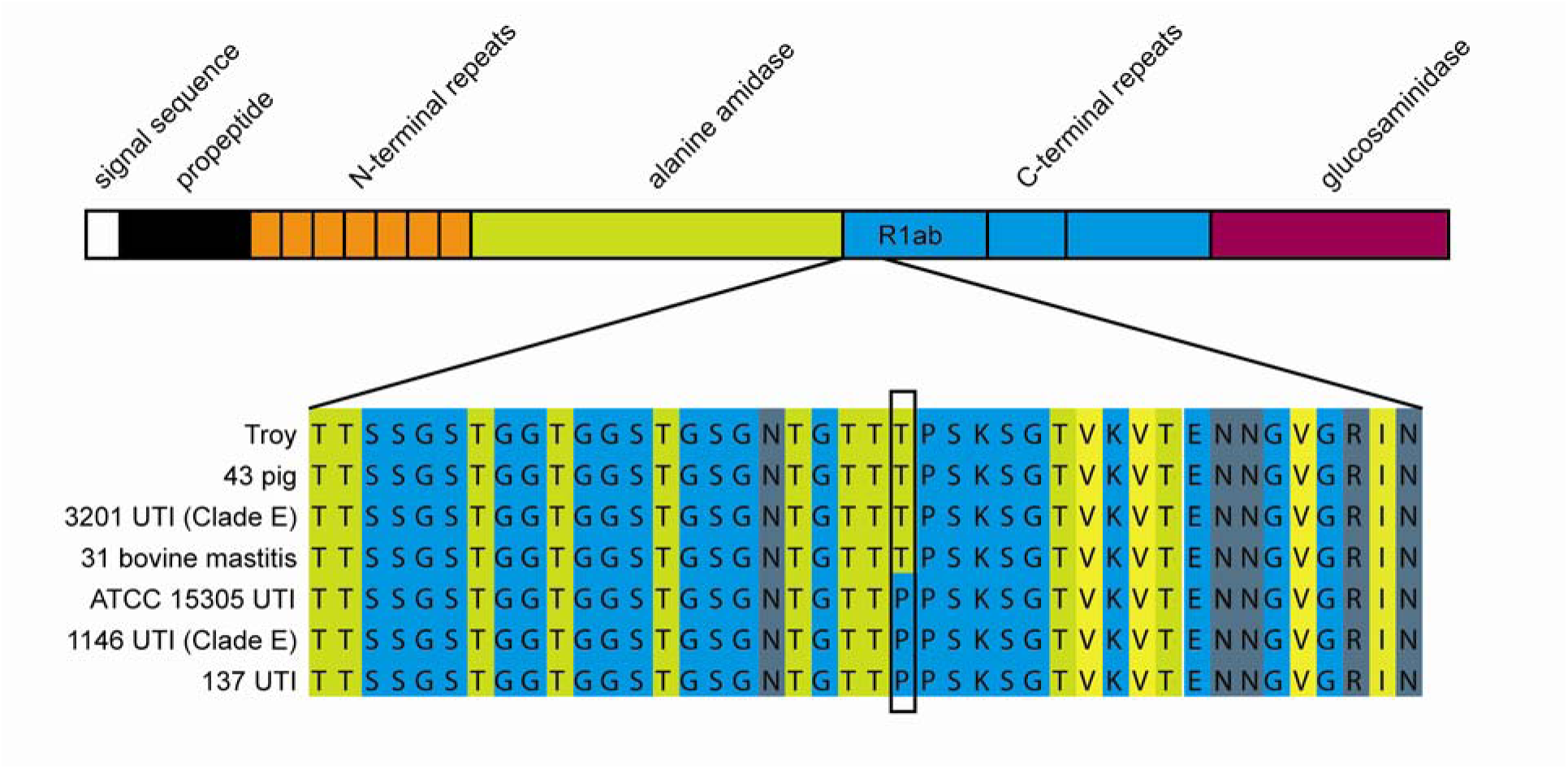
Non-synonymous variant in Aas fibronectin binding repeat. Top-Domains of Aas protein adapted from Hell et al. 1998. R1ab is the peptide used in the fibronectin and thrombospondin binding experiments. Bottom-Alignment of a portion of R1 showing amino acid sequence in Aas from selected *S. saprophyticus* strains. Amino acids are colored based on their propensity to form beta strands (light green=high propensity, light blue=low propensity). The alignment visualization was created in JalView.

The Aas variant is in a region known to bind fibronectin (Figure 5, (31)) and may be under selection because it affects adhesion to this host protein. We used ELISAs to investigate potential effects of *aas*_2206A>C on binding to fibronectin and thrombospondin-1, which binds to this region of the homologous AtlE amidase from *Staphylococcus epidermidis* (33). Staphylococcal autolysins contains 3 C-terminal repeats (R1-R3), which can each be divided into two subunits (a and b) based on structural information (34). We confirmed that Aas R1a1b binds fibronectin and discovered that it also binds thrombospondin (Figure 6); there was no detectable difference between the ancestral and derived R1a1b alleles in binding to fibronectin or thrombospondin (human or bovine).

**Figure 6.**
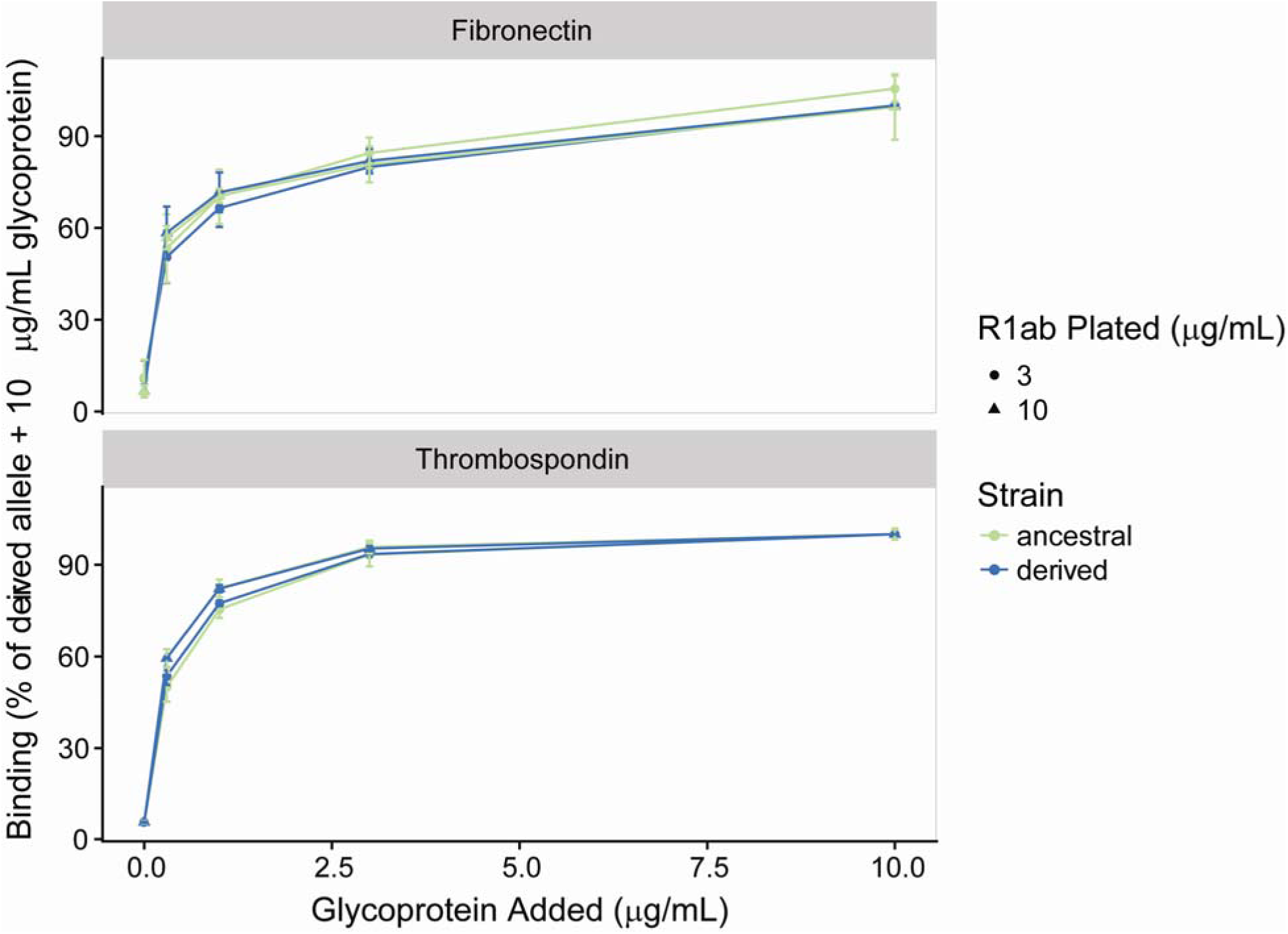
Fibronectin and thrombospondin binding to human-associated and ancestral strain Aas R1ab. ELISAs detecting the binding of soluble human fibronectin and thrombospondin to plates coated with Aas R1ab at 3 and 1 ug/ml. Results normalized to percent of binding of 10ug/ml glycoprotein to human-associated strain R1ab. Human and bovine fibronectin and human thrombospondin bound to the two constructs equally well.

Interestingly, we observed several instances of recombination of the *aas* variant. In each case, the recombination event reinforced the association of the derived allele with human infection. Two of the non-human-associated bacterial isolates in lineage U – an isolate from a pig and a second from cheese rind – had evidence of a recombination event at the *aas* locus resulting in acquisition of the ancestral allele. Conversely, one of the human UTI isolates in Clade E (for which the ancestral allele is otherwise fixed) acquired the derived *aas* variant.

Several human pathogens appear to have undergone recent population expansion (35–38). We wondered whether the uropathogenic lineage of *S. saprophyticus* might also have undergone a recent change in its effective size. The genome wide estimate of TD for lineage U was negative (-0.58), which is consistent with population expansion. We used the methods implemented in ∂a∂i (39) to identify the demographic model that best fit the observed synonymous SFS of lineage U (Figure 7).

**Figure 7.**
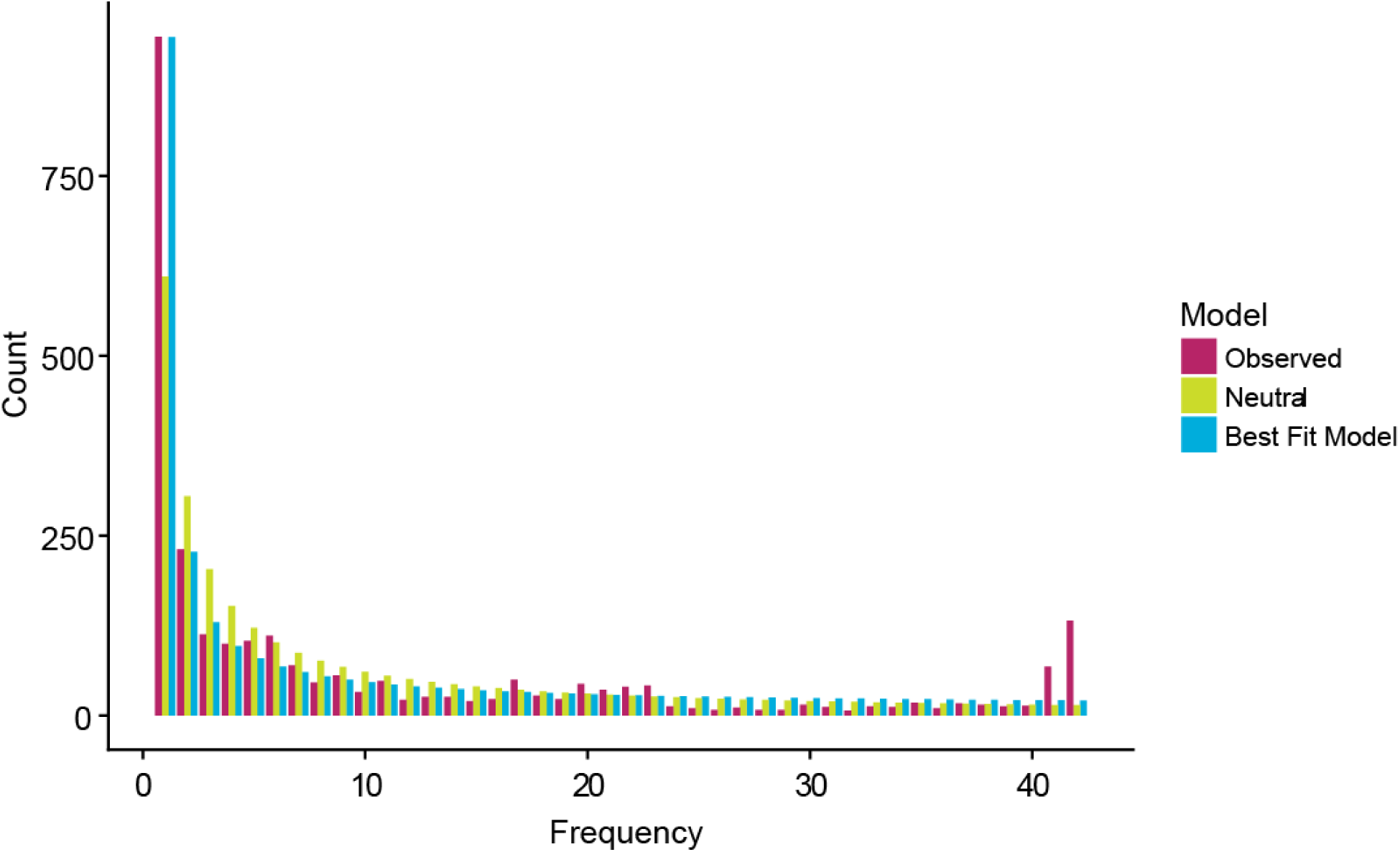
Site frequency spectrum of lineage U. The ancient genome (Troy) was used as the outgroup to determine the ancestral state. Synonymous, nonsynonymous, and intergenic sites were identified with SnpEff (98). The observed synonymous SFS contain an excess of singletons and high frequency derived variants. Both the observed SFS and the SFS predicted by the best fitting model have an excess of singletons compared to the SFS expected under the standard neutral model with no population size change.

The synonymous SFS showed an unexpected excess of high frequency derived alleles, which we hypothesized were the result of gene flow from populations with ancestral variants. Within population recombination has been shown to have no effect on SFS-based methods of demographic inference in bacteria (40). However, external sources of recombination were not modeled in previous studies. We used SimBac (41) to simulate bacterial populations with a range of internal and external recombination rates. Similar to previous studies, we did not find that within population recombination had an effect the value of Tajima’s D. However, we found that recombinant tracts from external sources resulted in positive values of Tajima’s D (Figure 8). Positive values of Tajima’s D are also associated with population bottlenecks and balancing selection.

**Figure 8.**
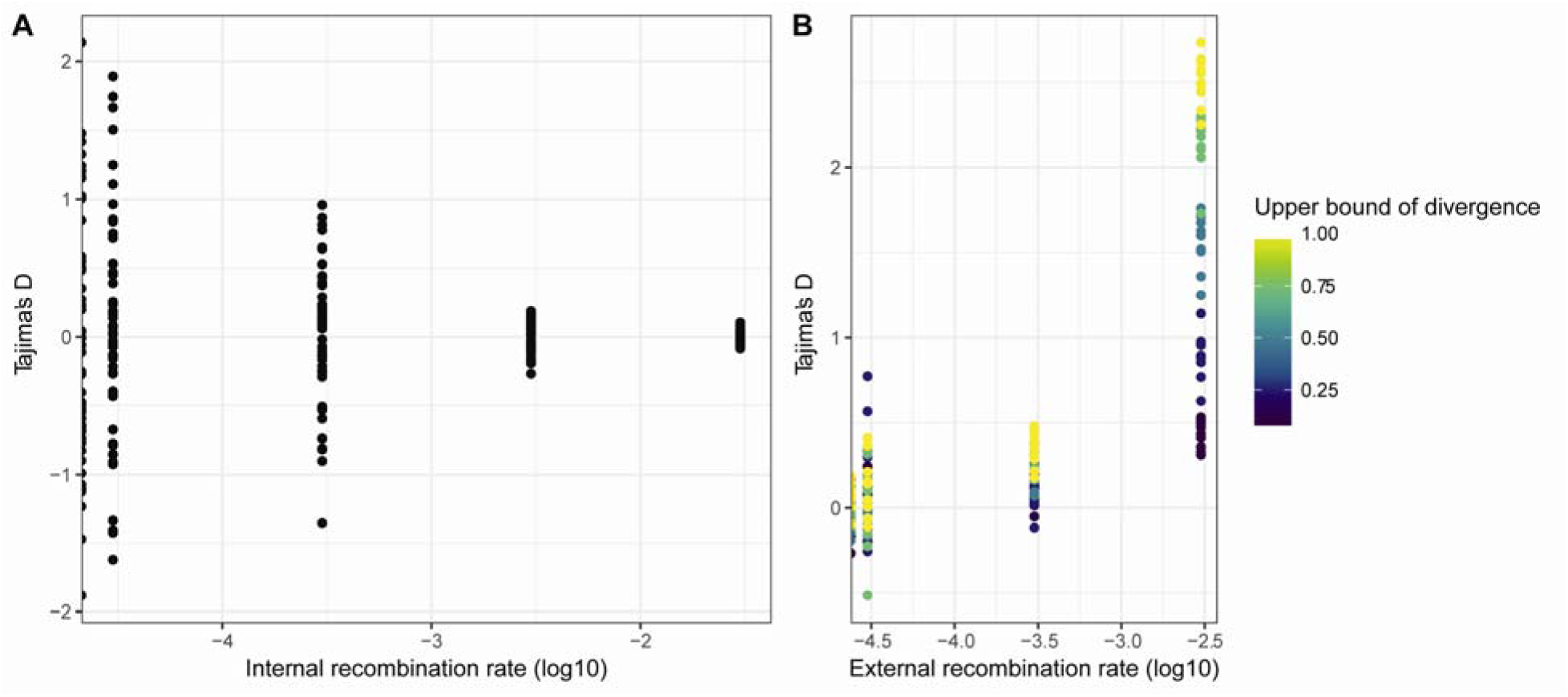
Effects of internal and external recombination on Tajima’s D. Bacterial populations with a range of recombination rates were simulated with SimBac. A) Taijma’s D values from simulations of internal recombination rates ranging from 0-0.03 and no external recombination. B) Tajima’s D values from simulations with an internal recombination rate of 0.003 (r/m = 1) and external recombination rates ranging from 0-0.003. Points are filled according to the upper limit of diversity in external recombinant fragments.

We used fastGEAR (42) to identify recombinant tracts that originated outside of lineage U, and these sites were removed from the analysis prior to demographic inference. We compared five demographic models (constant size, instantaneous population size change, exponential population size change, instantaneous population size change followed by exponential, and two instantaneous population size changes, Figure 9) and used bootstrapping to estimate the uncertainty of the parameters and to adjust the composite likelihoods using the Godambe Information Matrix implemented in ∂a∂i (43). We found significant evidence for an expansion in all models (Table 2). The best fitting model was an instantaneous contraction followed by an instantaneous expansion, in which the population underwent a tight bottleneck followed by a 15-fold expansion without recovering to its ancestral size (V = *N_e_*/*N_anc_*, T = generations/*N_anc_*, V_A_: 2.9 × 10^−2^, V_B_: 4.5 × 10^−1^, T_A_: 1.2 × 10^−1^, T_B_: 3.1 × 10^−3^).

**Figure 9.**
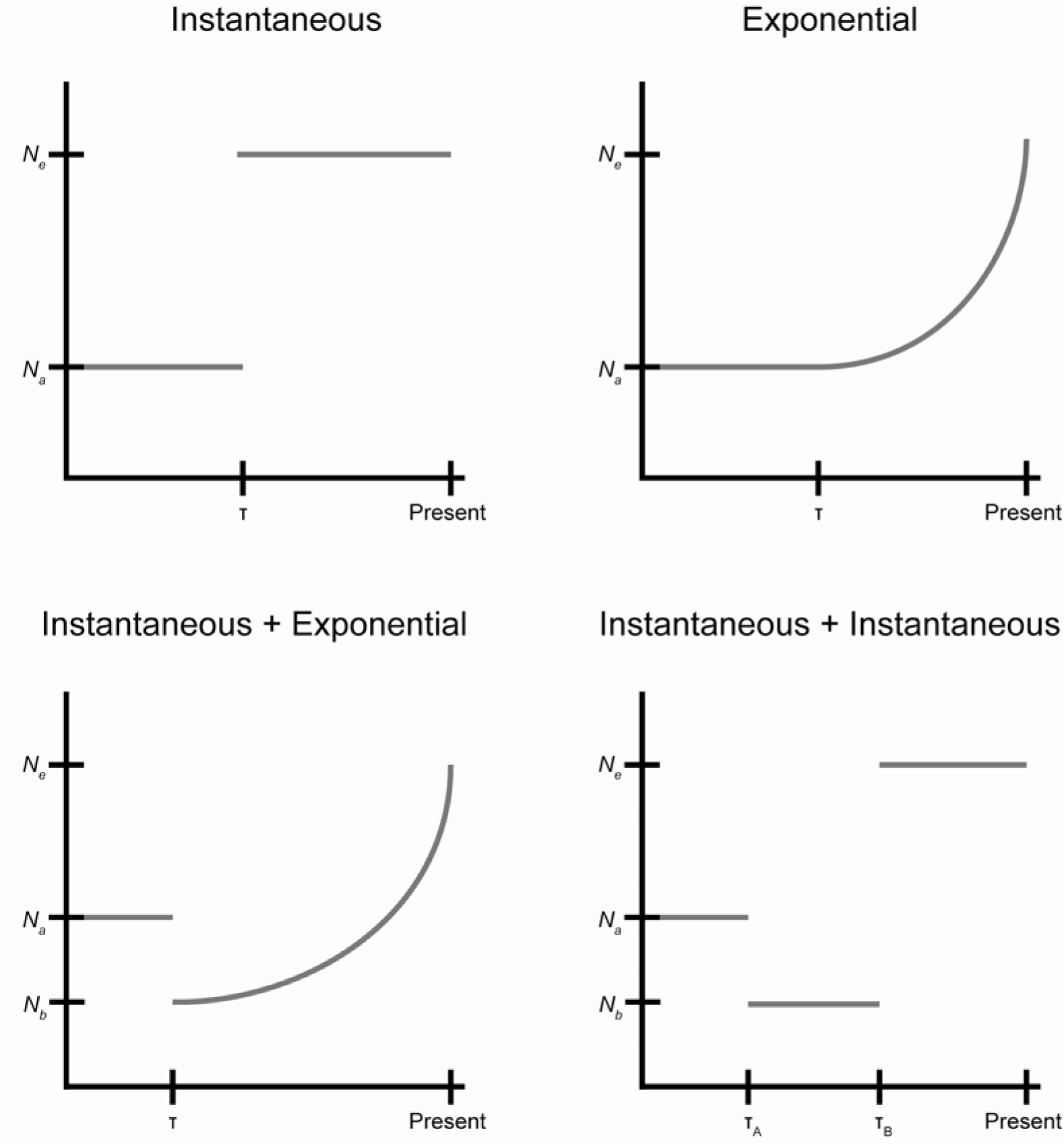
Cartoon of fitted demographic models. The observed synonymous SFS was fit to 5 demographic models including constant size, instantaneous population size change, exponential population size change, instantaneous population size change followed by exponential, and two instantaneous population size changes. Parameters for the instantaneous and exponential models are the magnitude of the population size change (V = *N*_*e*_/*N*_*ancestral*_) and the timing of the change (t = generations/*N*_*ancestral*_). For models with two population size changes, magnitudes are reported as V_A_ = *N*_*b*_/*N*_*ancestral*_ and V_b_ = *N*^e^/*N*^*ancestral*^.

**Table 2.** Results of demographic inference. V = *N*_*e*_/*N*_*ancestral*_, T = generations/*N*_*ancestral*_

| Model | Optimized parameters (standard deviation) | Log Likelihood | p-value (comparison model) |
| --- | --- | --- | --- |
| constant size |  | -562 |  |
| instantaneous change | $v: 9.9 \times 10^4 (6.0 \times 10^4)$<br>$\tau: 4.4 \times 10^{-2} (4.7 \times 10^{-3})$ | -455 | 0.0 (constant size) |
| exponential change | $v: 9.7 \times 10^4 (9.8 \times 10^4)$<br>$\tau: 4.1 \times 10^{-2} (4.9 \times 10^{-3})$ | -455 | 0.0 (constant size) |
| instantaneous change followed by exponential change | $v_A: 2.1 \times 10^{-2} (2.9 \times 10^{-2})$<br>$v_B: 1.1 (3.9 \times 10^{-2})$<br>$\tau: 1.3 \times 10^{-1} (1.6 \times 10^{-1})$ | -404 | $1.6 \times 10^{-4}$ (exponential change) |
| two instantaneous size changes | $v_A: 2.9 \times 10^{-2} (1.2 \times 10^{-2})$<br>$v_B: 4.5 \times 10^{-1} (2.1 \times 10^{-1})$<br>$\tau_A: 1.2 \times 10^{-1} (3.4 \times 10^{-2})$<br>$\tau_B: 3.1 \times 10^{-3} (5.3 \times 10^{-3})$ | -393 | $4.6 \times 10^{-7}$ (instantaneous change) |

Recombination and positive selection are known to confound the inference of bacterial demography (40), so we used simulations to investigate their effects on our demographic inference for uropathogenic *S. saprophyticus.* We used SFS_CODE (44) to simulate positive selection (with a range of recombination rates) and evaluate its effects on the accuracy of demographic inference with ∂a∂i. The method implemented in ∂a∂i relies on inference from the synonymous SFS, but it’s possible for synonymous variation to be affected by selection, particularly at low rates of recombination (40, 45). Neutral simulations with gene conversion did not affect demographic inference. We did find that positive selection can affect the synonymous SFS, resulting in inference of population size changes. In simulations of positive selection in a population of constant size, we found the spurious inference to be a bottleneck rather than an expansion. This suggests that the observed synonymous SFS of lineage U has been affected both by positive selection and by demographic expansion.

## Discussion

A central question in the population biology of infectious diseases is how and why pathogenic traits emerge in microbes. Addressing this question is important for understanding novel disease emergence and for identifying the genetic basis of virulence. Here we present evidence suggesting that a mutation in *S. saprophyticus*’ *aas*, which binds host matrix proteins, is under positive selection and has enabled emergence and spread of a human pathogenic, UTI-associated lineage of this bacterium.

*S. saprophyticus* is familiar to medical microbiologists and clinicians as a common cause of UTIs (46), which are associated with significant morbidity, economic costs, and severe complications (4). Despite its strong association with UTIs in humans, *S. saprophyticus* can also be isolated from diverse environments including livestock, food and food processing plants, and the environment (47, 48). Our previous research suggested that pathogenicity to humans is a derived trait in the species (7).

This pattern is replicated here, where phylogenetic analyses link human UTI with two lineages of *S. saprophyticus* that are nested among isolates from diverse, non-human niches (i.e. free living, food- and animal-associated). The *aas* mutation arose in lineage U, which contains most of the UTI isolates. Two lineages are basal to lineage U: one is bovine-associated, and the other contains an ancient bacterial sequence from a pregnancy-related infection in Late Byzantine Troy. The Troy bacterium has the ancestral, bovine-associated *aas* allele, and we have previously hypothesized (7) that this lineage could be associated with human infections in regions where humans have close contacts – e.g. shared living quarters– with livestock, as they did at Troy during this time.

A second cluster of UTI isolates appears in Clade E. One isolate has acquired the derived *aas* allele, which parallels our finding that two non-human isolates in lineage U acquired the ancestral variant; all recombination events that we observed at this locus reinforced the association between *aas*_2206A>C and human infection.

Several UTI isolates in Clade E do not have the derived *aas* allele and the clustering of UTI isolates suggests there may be a distinct adaptive path to virulence in this clade. Larger and more comprehensive samples will be needed to investigate this hypothesis and to identify the factors shaping the separation of clades P & E.

The *aas* mutation has characteristics associated with a classical selective sweep driven by positive selection, namely a regional reduction in diversity (21) and Tajima’s D (22, 49). With the exception of the interesting allelic replacements noted above, there was also relatively little recombination at this locus, consistent with it being functionally important. To our knowledge, this is the first description of a single nucleotide sweep in a bacterium.

Depending on the strength of selection and recombination rate, positive selection in bacteria has been observed to affect the entire genome, resulting in clonal replacements, or to only affect specific regions of the genome (50). For example, multiple clonal replacements have occurred in *Shigella sonnei* populations in Vietnam due to acquisition of resistance to antimicrobials and environmental stress (51). Recurrent clonal replacements have also been observed within single hosts during chronic infection of cystic fibrosis patients by *Pseudomonas aeruginosa* (52).

Environmental bacterial populations can also be subject to clonal replacements: a metagenomic time course study of Trout Bog found evidence of clonal replacement occurring in natural bacterial populations but not gene or region specific sweeps (53). However, large regions of low diversity were also observed, suggesting gene-specific selective sweeps had occurred prior to the start of the study. Shapiro et al. identified genomic loci that differentiated *Vibrio cyclitrophicus* associated with distinct niches but that had limited diversity within niches; they concluded that differentiation of these populations had been enabled by recombination events that reinforced the association of alleles with the niche in which they were advantageous (54).

The *aas*_2206A>C mutation is within a group of genetic variants that differentiates bacteria associated with human-pathogenic versus other niches (i.e. F_ST_ outlier). SNPs associated with specific clinical phenotypes were described recently in the pathogen *Streptococcus pyogenes* (55), which is consistent with our finding that clinical phenotypes can represent distinct niche spaces preferentially occupied by sub-populations of bacteria. There is also precedent for a single nucleotide polymorphism to affect host tropism of bacteria (56).

In sexually reproducing organisms, haplotype based statistics are frequently used to identify selective sweeps because positively selected alleles will also increase the frequency of nearby linked loci faster than recombination can disrupt linkage, producing longer haplotypes for selected alleles (28, 57). We found that *aas*_2206A>C had a longer haplotype than the ancestral variant, but this difference was not extreme relative to other regions of the genome (assessed with the *nS*_L_ statistic). Haplotype based statistics have been found to perform poorly in purebred dogs, where linkage across the genome is high (58). Relatively low levels of recombination may also contribute to a lack of sensitivity when haplotype-based detection methods are applied to bacteria; linkage of sites is also likely to be disrupted in a less predictable way by bacterial gene conversion than by crossing over (29). Based on our findings, we conclude that screening for regional decreases in diversity and distortions of the SFS (i.e. sliding window analyses) and identification of genetic variants with extreme differences in frequency between niches can be useful in identifying candidate sites of positive selection in bacteria.

*S. saprophyticus* encodes a number of adhesins including UafA, UafB, SdrI, and Aas. UafA and Aas are found in all isolates, suggesting that they play important roles in the diverse niches occupied by *S. saprophyticus*. Aas has autolytic, fibronectin binding, and haemagluttinating functions (9, 31, 32, 59). We identified a single, non-synonymous polymorphism as a target of selection in the fibronectin binding repeats of *Aas*. This variant is predicted to affect the repeat’s structure, as proline has a more rigid structure than other amino acids. Adhesins are plausible candidates for adaptation to the uropathogenic niche, as they are known to be important virulence factors in pathogens causing urinary tract infections (60). Fibronectin binding proteins including Aas have been identified as virulence factors in *S. saprophyticus* and *Enterococcus faecalis* (32, 61, 62). Adhesion to the uroepithelium is essential for uropathogens to establish themselves in the bladder, where they are subject to strong shear stress (63): we hypothesize that *S. saprophyticus* with the derived *aas* variant are better able to colonize the human bladder.

Invasion of the human urinary tract may provide a fitness advantage by allowing relative enrichment of *S. saprophyticus* in a site with little competition from other bacterial species and by providing a mechanism of dispersal in the environment. In analyses of selection in *Escherichia coli*, another bacterium occupying diverse niches, residues in the adhesin FimH were found to be subject to positive selection in uropathogenic strains (64–66). FimH binds mannose, providing protection from shear stress through a catch bond mechanism (67). Interestingly, *Borrelia burgdorferi*’s vascular adherence and resistance to shear stress were recently found to be enabled by interactions between a bacterial adhesin and host fibronectin that also use a catch bond mechanism (68). There are also precedents in *Staphylococcus aureus* for polymorphisms in bacterial fibronectin-binding adhesins to affect the strength of binding, and for these polymorphisms to associate with specific clinical phenotypes (69).

Further experiments are needed to investigate the effects of variation in Aas on *S. saprophyticus* biology. In our preliminary investigations of binding using ELISA assays of recombinant bacterial peptides, we did not detect differences between ancestral and derived alleles in binding of the R1a1b repeat to fibronectin. The variant could still affect fibronectin binding by altering conformation of the protein in a manner analogous to FimH in *E. coli* (66). It’s also possible that variants in the peptide affect binding under specific conditions that we did not test. Another possibility is that the variant affects autolysis or other as yet undescribed functions of Aas. The roles of adhesins and other virulence factors in *S. saprophyticus*’s colonization of niches in livestock and the environment are also interesting topics for further study.

Our demographic analysis of the uropathogenic lineage of *S. saprophyticus* showed evidence of a population bottleneck and subsequent expansion. Bottlenecks and expansion of drug resistant clones have previously been shown to affect the population structure of *Streptococcus agalactiae* (70), demonstrating the effects of positive selection on the demographic trajectories of bacterial sub-populations. However, previous work has also shown that selection - and recombination - can produce spurious results from demographic inference in bacteria (40, 71). We used an SFS-based method to reconstruct the demographic history of *S. saprophyticus*: the accuracy of demographic inference using these methods has been shown to be unaffected by within-population recombination (40) and this was confirmed in our analyses of simulated data. We found that recombination from external sources may result in an excess of intermediate frequency variants, which is also a signature of population bottlenecks, so we masked externally imported sites. However, the frequency of synonymous variants could still be affected by selection on linked non-synonymous sites, including the selective sweep in *aas* that we have described. We performed simulations to address these potential confounders and aid in the interpretation of our demographic inferences. Simulation of a single site under positive selection resulted in the inference of a bottleneck (*N*_*e*_/*N*_*a*_ = 0.01-0.42), indicating that, at the recombination rates we simulated, diversity was lost from neutrally evolving sites due to their linkage to the site under selection. In inferences from our observed data, a bottleneck was followed by a 15-fold expansion, suggesting that lineage U has undergone both a selective sweep and demographic expansion.

Here we have described adaptation of *S. saprophyticus* that may have enabled its expansion into a human pathogenic niche. Mutation of a single nucleotide within the *aas* adhesin appears to have driven a selective sweep, and allele frequency differences at the locus are consistent with niche-specific adaptation. Lateral gene transfer events in *aas* reinforced the association of the positively selected allele with human infection. These results provide new insights into the emergence of virulence in bacteria and outline an approach for discovering the molecular basis of adaptation to the human pathogenic niche.

## Methods

### DNA extraction

After overnight growth in TSB at 37°C in a shaking incubator, cultures were pelleted and resuspended in 140 μL TE buffer. Cells were incubated overnight with 50 units of mutanolysin. We used the MasterPure Gram Positive DNA Purification Kit (EpiCentre) for DNA extraction.For DNA precipitation we used 1 mL 70% ethanol and centrigured at 4°C for 10 minutes. We additionally used a SpeedVac for 10 minutes to ensure pellets were dry before re-suspending the pellet in 50 µL water.

### Library preparation and sequencing

For SSC01, SSC02, and SSC03, library prep was performed using a modified Nextera protocol as decribed by Baym et al. (72) with a reconditioning PCR with fresh primers and polymerase for an additional 5 PCR cycles to minimize chimeras and a two-step bead based size selection with target fragment size of 650 bp and sequenced on an Illumina HiSeq 2500 (paired-end, 150 bp). For 43, SSC04, SCC05, and SSMast, DNA was submitted to the University of Wisconsin-Madison Biotechnology Center for library preparation and were prepared according the TruSeq Nano DNA LT Library Prep Kit (Illumina Inc., San Diego, California, USA) with minor modifications. A maximum of 200 ng of each sample was sheared using a Covaris M220 Ultrasonicator (Covaris Inc, Woburn, MA, USA). Sheared samples were size selected for an average insert size of 550 bp using Spri bead based size exclusion. Quality and quantity of the finished libraries were assessed using an Agilent DNA High Sensitivity chip (Agilent Technologies, Santa Clara, CA) and Qubit dsDNA HS Assay Kit, respectively. Libraries were standardized to 2 μM. Paired-end, 150 bp sequencing was performed using v2 SBS chemistry on an Illumina MiSeq sequencer. Images were analyzed using the Illumina Pipeline, version 1.8.2.

### Reference guided mapping

We mapped reads to ATCC 15305 using via a pipeline (available at https://github.com/pepperell-lab/RGAPepPipe). Briefly, read quality was assessed and trimmed with TrimGalore! v 0.4.0 (www.bioinformatics.babraham.ac.uk/projects/trim_galore), which runs both FastQC (www.bioinformatics.babraham.ac.uk/projects/fastqc) and cutadapt. Reads were mapped to using BWA-MEM v 0.7.12 (73) and sorted using Samtools v 1.2 (74). We used Picard v 1.138 (picard.sourceforge.net) to add read group information and removed duplicates. Reads were locally realigned using GATK v 2.8.1 (75). We identified variants using Pilon v 1.16 (76) (minimum read depth: 10, minimum mapping quality: 40, minimum base quality: 20).

### Assembly

We used the iMetAMOS pipeline for *de novo* assembly (77). We chose to compare assemblies from SPAdes (78), MaSurCA (79), and Velvet (80). KmerGenie (81) was used to select kmer sizes for assembly. iMetAMOS uses FastQC, QUAST (82), REAPR (83), LAP (84), ALE (85), FreeBayes (86), and CGAL (87) to evaluate the quality of reads and assemblies. We also used Kraken (88) to detect potential contamination in sequence data. For all newly assembled isolates (43, SSC01-05, SSMast), the SPAdes assembly was the highest quality. Assembly statistics are reported in Table S2.

### Annotation and gene content analyses

We annotated the de novo assemblies using Prokka v 1.11 (89) and used Roary (90) to identify orthologous genes in the core and accessory genomes. To look for associations between accessory gene content and human association, we used Scoary (19). For the analysis, we used human association as our trait, and we adjusted the p-value for multiple comparisons using the Bonferroni method.

### Alignment

When short read data for reference guided mapping were unavailable, whole genome alignment of genomes to ATCC 15305 was performed using Mugsy v 2.3 (91). Repetitive regions in the reference genome greater than 100 bp were identified using nucmer, and these regions were masked in the alignment used in downstream analyses.

### Maximum likelihood phylogenetic analysis

Maximum likelihood phylogenetic trees were inferred using RAxML 8.0.6 (92). We used the GTRGAMMA substitution model and performed bootstrapping using the autoMR convergence criteria. Tree visualizations were created in ggtree (93).

### Population genetics statistics

To calculate π and Tajima’s D, we used EggLib v 2.1.10 (94), a Python package for population genetic analyses. A script to perform the sliding window analysis is available at https://github.com/tatumdmortimer/popgen-stats/slidingWindowStats.py. We used vcflib (https://github.com/vcflib/vcflib) to calculate F_ST_ and EHH and selscan v 1.1.0b (95) to calculate *nS*_L_.

### Recombination analyses

To identify recombinant regions in the *S. saprophyticus* alignment, we used Gubbins v 2.1.0 (20). fastGEAR (42) was used with the recommended input specifications to identify recombination events between major lineages of *S. saprophyticus*. We used Circos (96) for visualization of recombinant tracts.

### Site frequency spectrum

We used SNP-sites v 2.0.3 (97) to convert the alignment of *S. saprophyticus* isolates to a multi-sample VCF. SnpEff v 4.1j (98) was used to annotate variants in this VCF as synonymous, non-synonymous, or intergenic. Using the Troy genome as an outgroup, we calculated an unfolded site frequency spectrum (SFS) for lineage U for each category of sites. To reduce the impact of lateral gene transfer on the SFS, we removed sites where the origin was outside lineage U based on results of fastGEAR.

### *Ancestral reconstruction of aas*_2206A>C

We used TreeTime (99) to reconstruct the evolutionary history of the variant in *aas* using the maximum likelihood phylogeny inferred using RAxML.

### Demography

We performed demographic inference with the synonymous SFS using ∂a∂i (39). Models tested were the standard neutral model, expansion, and exponential growth. Parameters, V (*N_e_*/*N_a_*) and T (time scaled by 2), were optimized for both the expansion and exponential growth models. Significance of the expansion and exponential growth model compared to the standard neutral model was evaluated using a likelihood ratio test. The scripts used to perform this analysis are available at https://github.com/tatumdmortimer/popgen-stats.

### Simulations

Simulations were performed in SimBac (41) to evaluate the effect of external recombination on the SFS. Populations were simulated with sample size and θ equivalent to our sample of lineage U (*n* = 44, θ = 0.003). The length of internal recombinant tracts was 6500 bp (median of Gubbins output), and the length of external recombination events was 3000 bp (median of fastGEAR output). Internal recombination was simulated at rates ranging from 0 to 0.03 (r/m = 10). External recombination was simulated at rates ranging from 0 to 0.003. The lower bound of difference for external recombination was 0, and the upper bound was simulated at ranges from 0.25-1.0. Simulations were performed in SFS_CODE (release date 9/10/2015) (44) to evaluate the power of ∂a∂i to accurately estimate demographic parameters in the presence of gene conversion and selection. We simulated a locus of length 100 kb with theta 0.003, gene conversion tract length of 1345 bp, and a range of recombination/mutation ratios (0.0002-2.0). In addition to neutral simulations with gene conversion, we also performed simulations with a single site under selection (γ = 10-1000) with the same parameters as neutral simulations.

### Expression of R1ab

Human-associated and ancestral strain R1ab were cloned into the expression vector pET-ELMER (LM Maurer 2010, JBC), transformed into BL21(DE3)cells (EMD, Gibbstown, NJ) for expression and induced with 1mM IPTG. Bacteria were lysed in 100 mM NaH_2_PO_4_, 10 mM Tris, 8M urea, 1 mM β-mercapto-ethanol, 5 mM imidazole, pH 8.0 (lysis buffer). The cleared lysate was incubated overnight with nickel-nitrilotriacetic acid agarose (Qiagen), washed and eluted in lysis buffer pH 7.0 plus 300 mM imidazole.

### ELISA

Antigen was diluted to 10 µg/ml in TBS (10 mM Tris, 150 mM, pH 7.4) and used to coat 96-well microtiter plates (Costar 3590 high binding, Corning Inc., Corning, NY) with 50 µl per well, for 16 h at 4□C. The plates were blocked with 1% BSA in TBS plus 0.05% Tween-20 (TBST) for 1 h. After washing three times with TBST, purified plasma fibronectin (100) or platelet-derived thrombospondin-1 (101) diluted to 10, 3, 1, or 0.3µg/ml in TBST plus 0.1% BSA, were added to the plates and incubated for 2 h. Plates were washed four times with TBST. Rabbit anti-fibronectin and rabbit anti-thrombospondin antibodies diluted in TBST plus 0.1% BSA were added to the appropriate wells and incubated for 1 h. Plates were washed four times with TBST. Peroxidase-conjugated secondary antibody was incubated with the plates for 1 h. Plates were washed four times with TBST and 50 µl per well of SureBlue TMB peroxidase substrate (KLP) was added to each well. Color development was monitored for 10-30 min, 50 µl of TMB stop solution (KLP) was added, followed by measurement of absorbance at 450nm.

## Acknowledgments

We thank Andrew Kitchen (University of Iowa), Jeniel Nett (UW-Madison), JD Sauer (UW-Madison), and Rod Welch (UW-Madison) for their helpful input on this study. We also thank the University of Wisconsin Biotechnology Center DNA Sequencing Facility for providing library preparation and sequencing facilities and services.

## Supplementary Tables

**Table S1.** Accession numbers for *S. saprophyticus* isolates.

**Table S2.** Assembly statistics for *S. saprophyticus* genomes.

